# Evolution of functional genomic diversity during a bottleneck

**DOI:** 10.1101/2025.01.16.632721

**Authors:** Flávia Schlichta, Stephan Peischl, Laurent Excoffier

## Abstract

Most species have been through population bottlenecks and range expansions, and the impact of these events on patterns of diversity has been well studied. In particular, it has been shown that initially rare neutral variants could readily fix on the front of range expansions or during bottlenecks, giving genomic signatures looking like selective sweeps. Here we expand on previous work by considering the dynamics of genomic diversity in (functional) regions harbouring deleterious variants during bottlenecks or during range expansions modelled as serial founder effects. We find that regions with very low levels of diversity (troughs) looking like selective sweeps can also readily form in these functional regions. Additionally, their properties depend on the dominance level of deleterious mutations. Trough density is higher and increases more rapidly in regions with co-dominant deleterious mutations than in regions with recessive mutations. Interestingly, we find that genetic diversity declines less rapidly in regions with partially recessive mutations than in regions with codominant ones or in regions with only neutral mutations. These features are generally enhanced in low recombination regions and for intermediate selection coefficients. If most deleterious mutations in a genome are partially recessive, it follows that functional low recombination regions should better preserve genetic diversity during range expansions than neutral regions of the genome.

**Significance statement:** Most research on the effects of selection on genetic diversity have been done in equilibrium populations, leaving a gap in our understanding of how selection acting on functional regions interacts with demographic changes. Our study reveals strikingly different impacts of purifying selection on genetic diversity depending on the dominance coefficient of deleterious mutations. While codominant variants create diversity dips resembling selective sweeps, recessive mutations can preserve genetic diversity during bottlenecks through the relatively understudied phenomenon of pseudo-overdominance. Our findings show how selective and demographic forces interact in more complex ways than previously thought and emphasize the importance of further exploring these dynamics to accurately interpret genetic variation in natural populations.

## Introduction

Genetic variation is a product of the input of variants through mutation and the maintenance or removal of these variants by genetic drift or natural selection. Generally, selection is the driving force in large populations while drift becomes more important in smaller populations. Thus, to understand current levels of diversity, we must be able to disentangle these two competing forces. This is not an easy feat, however, since demographic changes can affect genomic diversity in ways similar to selection. For example, one of the earliest known signatures of positive selection is a decrease in heterozygosity around selected loci (Maynard-Smith and Haigh 1974; Fay and Wu 2005), but such localized depletions of diversity are also a common outcome in population having gone through bottlenecks (Moinet et al. 2022) or after range expansions (Schlichta et al. 2022). Also, population expansions generate a coalescent tree similar to those from selective sweeps, creating a skewed site frequency spectrum (SFS, (Wakeley 2009). Akin yet distinct, the slow purging of deleterious mutations (or background selection, BGS) distorts the SFS in similar ways by producing an excess of low frequency variants (Desai et al. 2012; Cvijović et al. 2018). Contrastingly, an excess of high frequency variants can be caused by both recent selective sweeps and bottlenecks (Wakeley 2021; Moinet et al. 2022). The effect of positive or negative selection on nearby diversity is also dependent on local levels of recombination, with more pronounced effects in regions of low recombination (Begun and Aquadro 1992; Charlesworth et al. 1993; Hudson and Kaplan 1995; Stephan 2010; Pouyet et al. 2018).

Due to all these confounding characteristics, much of the early work on selection has been done assuming stable populations at mutation-drift-equilibrium, but several researchers have pushed for the development of more complex “null-models” (Cutter and Payseur 2013; Bank et al. 2014; Kern and Hahn 2018; Johri et al. 2020; Smith and Hahn 2024), and trying to account for BGS in selection scans (Huber et al. 2016). Recently, researchers have also started to study the combined effects of BGS and changes in population size to obtain better expectations of their consequences but also to improve past demography inference (Torres et al. 2020; Johri et al. 2021). However, most studies investigating the action of selection are based on codominant variants (associated with dominance coefficient *h* = 0.5), which is not necessarily realistic (Di and Lohmueller 2024), since the fate of codominant and (partially) recessive selected variants can be quite different after demographic changes. For instance, after a bottleneck, deleterious recessive mutations are more easily purged, while codominant variants might actually increase in frequency (Kirkpatrick and Jarne 2000; Balick et al. 2015). Also, beneficial recessive variants spend longer times at low frequencies compared to dominant variants, thus having more opportunities to recombine into different backgrounds before reaching fixation, thus preserving more linked diversity during sweeps (Teshima and Przeworski 2006). Therefore, studies investigating the interaction between demographic changes and selection not only need to take into account the selection coefficients of mutations but also their levels of dominance.

Considering the importance of selection in general and of BGS in particular for shaping patterns of genomic diversity (McVean and Charlesworth 2000; Barton and Etheridge 2004; Stephan 2010; Charlesworth 2013; Pouyet et al. 2018; Gilbert et al. 2020) we aim here at expanding our previous work on the effect of range expansions on genomic diversity in neutral regions where we showed that regions of low diversity (troughs) would readily form during such expansions (Schlichta et al. 2022), and that their properties were resembling those of selective sweep (Moinet et al. 2022). In particular, we want to examine whether false selective sweep signals identified as troughs can still occur in functional regions after a series of bottlenecks, whether their dynamics is hampered or facilitated by BGS in regions harbouring a mixture of neutral and deleterious variants with contrasting dominance coefficients, and to what extent recombination can modulate these signals.

## Results

### A simple bottleneck mimics the effect of a range expansion

In a previous paper (Schlichta et al. 2022), we studied the effect of a range expansion on patterns of genomic diversity. In particular, we studied how the increase in frequency (and even fixation) of small chromosomal segments due to genetic surfing (Edmonds et al. 2004; Klopfstein et al. 2006; Hallatschek and Nelson 2008) generates patterns of low diversity on the front of a range expansion. Since range expansions can be modelled as a series of recurrent founder effects in a single population (DeGiorgio et al. 2011; Slatkin and Excoffier 2012), we compared patterns of diversity occurring during a bottleneck and during a range expansion (**supplementary fig. S1**). We found that conditional on the amount of diversity lost since the beginning of the bottleneck or the range expansion, bottlenecks of different sizes lead to virtually the same dynamics of trough formation than that observed during a range expansion.

This observation justifies the use of simple bottleneck simulations in the following rather than more complex spatially explicit range expansions and suggests that all results obtained from bottleneck simulations should also apply to what happens at the edge of spatial range expansions. Thus, we performed forward simulations of a large ancestral population (𝑁_𝑎𝑛𝑐_ = 10,000 diploid individuals) going through a sudden and severe bottleneck. To track the decay of diversity over time, we sampled the bottlenecked population at 5-generation intervals and generated genomic diversity scans to characterize chromosome segments of highly reduced diversity, hereafter referred as troughs. Troughs were defined as regions with 10% or less of the average diversity observed in the ancestral population. An illustration of the genomic scans showing the global changes in patterns of genomic diversity and of troughs formation over time is shown in **fig. 1**. Our simulation conditions were broadly based on human genomic diversity, see Material and Methods for further details.

**Figure 1:**
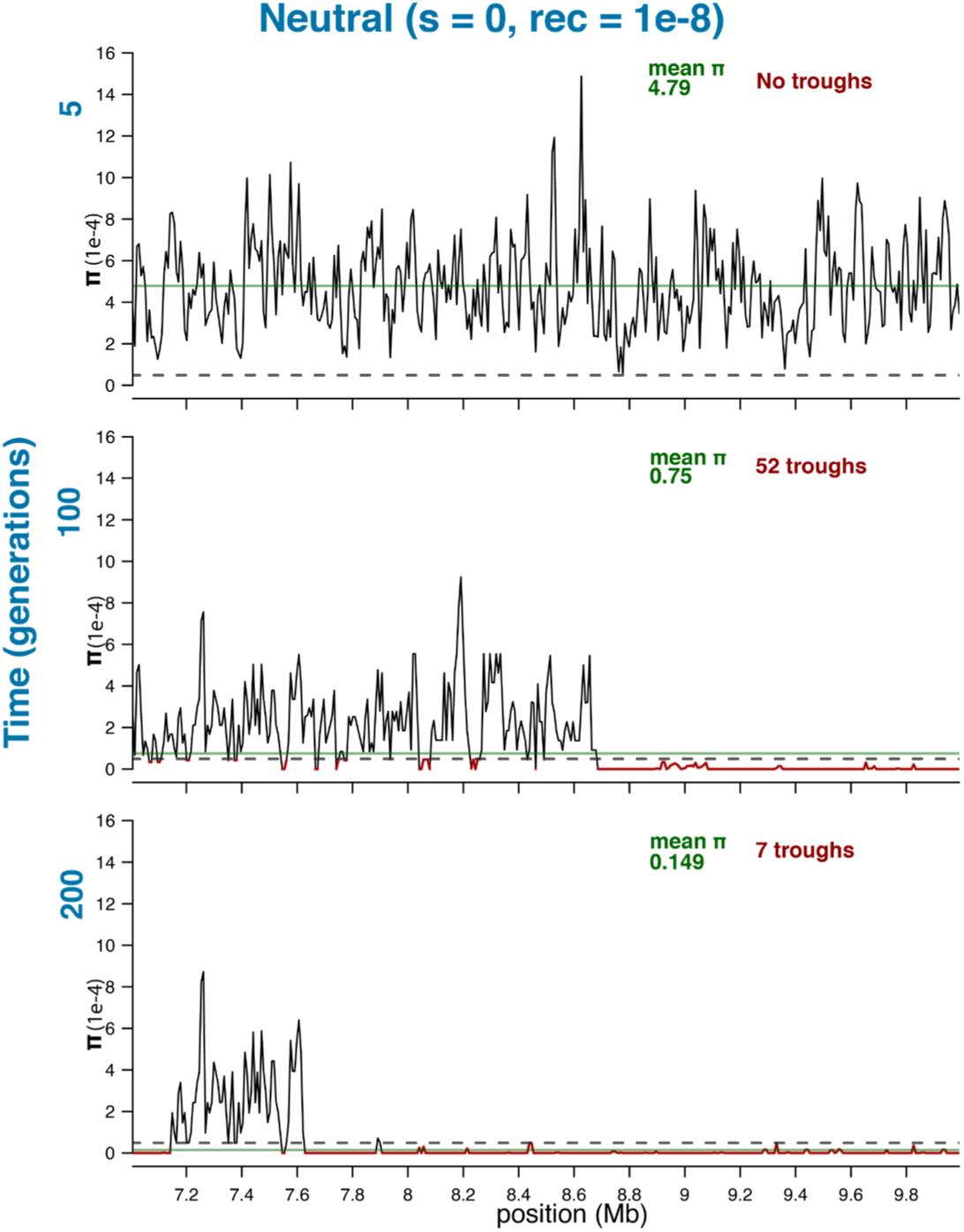
Genome scans of nucleotide diversity (π) during a bottleneck, for a neutral region of 3 Mb. Three time points are shown: 5, 100 and 200 generations after the start of the bottleneck. Y-axis shows nucleotide diversity and X-axis shows position in the genome. The horizontal dotted line indicates the threshold used to define throughs (10% of average ancestral diversity). The horizontal solid green line shows the average diversity of the chromosome segment at each three-time point, and the exact value is mentioned in green on the top of each panel. Troughs are highlighted in red, with their total numbers (for the 10Mb chromosome) shown in red on the top right of each panel. Details on the parameters used in the neutral forward simulations can be found in the Material and Methods section.

### Importance of ancestral population size in shaping diversity changes during a bottleneck

Since the effects of BGS on genomic diversity can be approximated as a local reduction in effective population size (Charlesworth et al. 1993; Nordborg et al. 1996; Charlesworth 2013), we compare trough formation during a bottleneck in populations with varying ancestral sizes. Despite differences in 𝑁_𝑎𝑛𝑐_, trough formation during a bottleneck of a given size (*N_Bot_*) leads to qualitatively very similar dynamic: following population size reduction, the trough density increases rapidly, reaches a maximum density at ∼ 2*N_Bot_* generations, and then decreases as troughs begin to merge, resulting in an increase in their average size (**fig. 2**). Quantitatively, however, the initial level of diversity does matter for trough dynamics.

**Figure 2:**
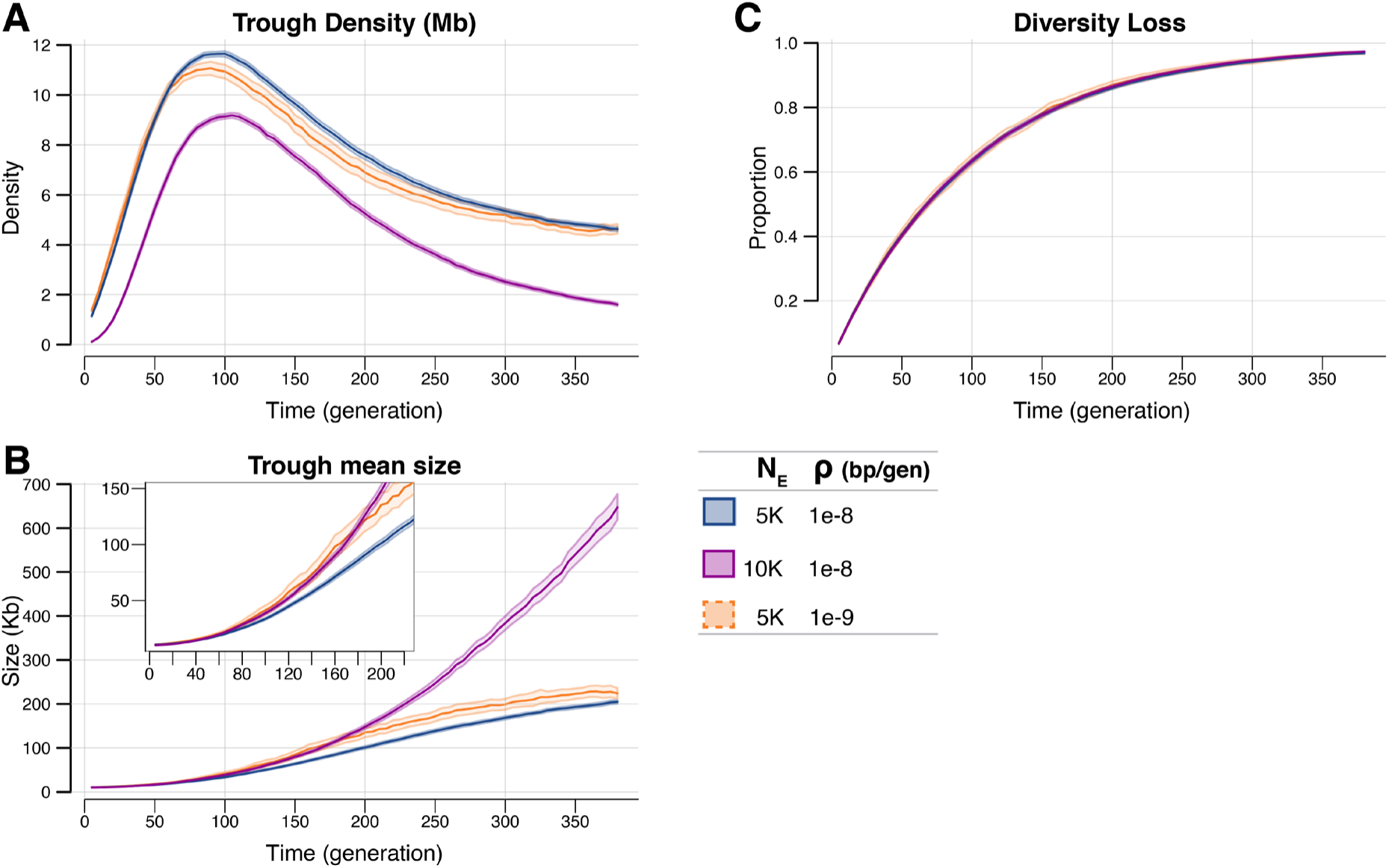
Statistics of trough dynamics during a bottleneck for neutral simulations in populations having different ancestral sizes (𝑁_𝑎𝑛𝑐_) or different recombination rates (𝝆). (A) Trough density (no. of troughs per Mb), (B) average trough size and (C) diversity loss relative to the ancestral population. Populations with smaller ancestral diversity (blue and orange lines) show higher trough density and smaller troughs compared to the population with larger ancestral sizes (magenta) before the onset of the bottleneck. Lower recombination (orange) only has a small effect in trough size and trough density. Importantly, despite different trough statistics and starting levels of diversity, all populations lose diversity at the same rate. Shaded areas show 95% CI obtained from the bootstrap distribution of 10,000 bootstrap samples corresponding to resampling 100 genomic simulations with repetition.

Indeed, we see that more troughs (**fig. 2A**) of smaller size (**fig. 2B**) are created when a population with a smaller ancestral size (with less standing genetic variability) goes through the same bottleneck as an initially larger population. However, the rate of diversity loss does not depend on the initial level of diversity as overall genetic diversity (average heterozygosity) diminishes identically when starting from a high or a low level of diversity (**fig. 2C**). Moreover, recombination rate slightly affects trough formation, with regions of low recombination showing comparatively larger but less numerous troughs than regions with higher recombination rates. But again, the overall change in diversity level does not depend on recombination rate.

### Effect of deleterious mutations in the ancestral population

We then simulated real BGS in the ancestral population by adding deleterious mutations at rate µ_𝐷𝑒𝑙_ = 1.1 ×10^-9^ in a portion of the genome with lower recombination rate 𝜌_𝐷𝑒𝑙_ = 5 × 10^-9^, since regions with recombination rate in humans higher than 1cM/Mb (1 × 10^-8^) are largely unaffected by BGS (Pouyet et al. 2018). Deleterious variants had a fixed selection coefficient *s* and identical dominance coefficient *h* for all variants (either highly recessive with h = 0.1, or codominant h = 0.5). We compare the diversity of BGS-affected regions to neutral regions using a common measure of background selection strength, *B*, which is defined as the ratio 𝜋_𝐵𝐺𝑆_/𝜋_𝑁𝑒𝑢_, where 𝜋 represents the average diversity in a given region. This metric provides a clear indication of how much diversity is reduced in BGS-affected regions relative to neutral regions. As expected, the presence of deleterious mutations over a genomic region affects the level of ancestral diversity, as reflected by the B-statistic shown in **Table 1** for various combinations of selection coefficients, dominance levels and recombination rates. For co-dominant deleterious mutations, the ancestral level of diversity in the selected chromosome is always lower than in the neutral chromosome, with much lower diversity with higher selection coefficients and in regions of low recombination, in line with expectations (Charlesworth et al. 1993; Hudson and Kaplan 1995). Contrastingly, in the presence of partially recessive variants, the level of diversity can be either lower or higher than that in the neutral chromosome. For the same recombination rate, diversity goes down with increasing selection coefficients. However, in regions of low recombination, nucleotide diversity can be much higher in the selected than in the neutral chromosome, especially in the presence of deleterious mutations with small selection coefficient (s = 0.0001). This result can be attributed to the presence of associative (pseudo-)overdominance (Sved 1968; Ohta and Kimura 1969; Ohta 1971; Zhao and Charlesworth 2016; Gilbert et al. 2020; Glémin 2022) and has been already observed in other simulations studies (Smith and Hahn 2024) and in real organisms (Schou et al. 2017). This transition from BGS to associative (pseudo-)overdominance is thus expected in regions of low recombination where recessive mutations at multiple loci are captured in repulsion on different haplotypes (Gilbert et al. 2020).

**Table 1:**
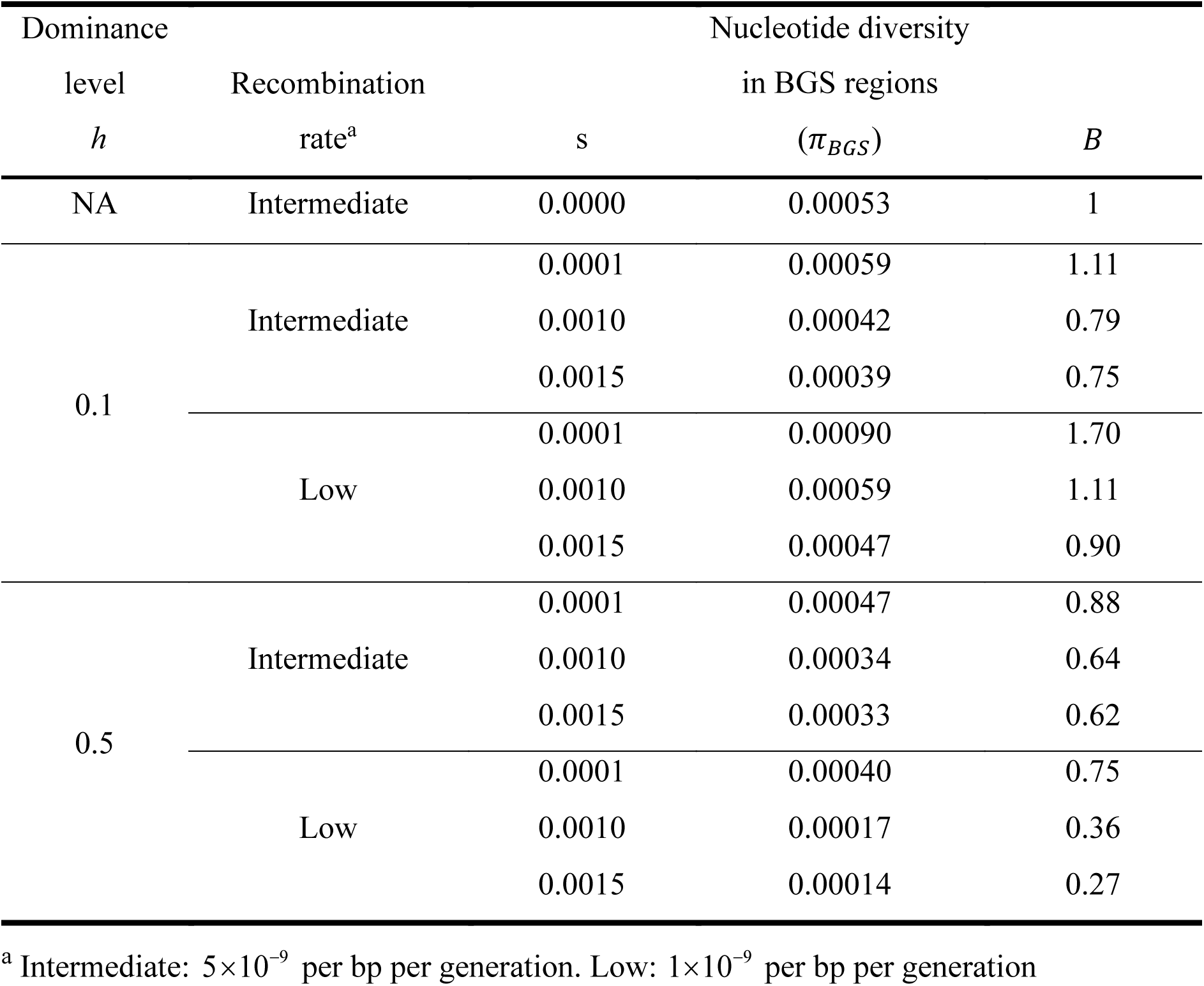
BGS intensity (measured by *B* values) for cases with different selection intensities (*s*), dominance coefficients (*h*) and recombination rates.

### Effect of deleterious mutations during the bottleneck

Whereas we have seen that levels of (neutral) ancestral diversity have a mild effect on trough properties during a bottleneck (**fig. 2A-B**), the rate at which diversity is lost is not affected by ancestral diversity (**fig. 2C**). Contrastingly, we find that in the presence of deleterious mutations, both trough dynamics and rate of diversity loss differ from what is expected under neutrality. In **fig. 3** we show the effect of dominance level and recombination rate on genomic diversity, whereas in **fig. 4** we show the effect of selection intensity and recombination for highly recessive mutations.

**Figure 3:**
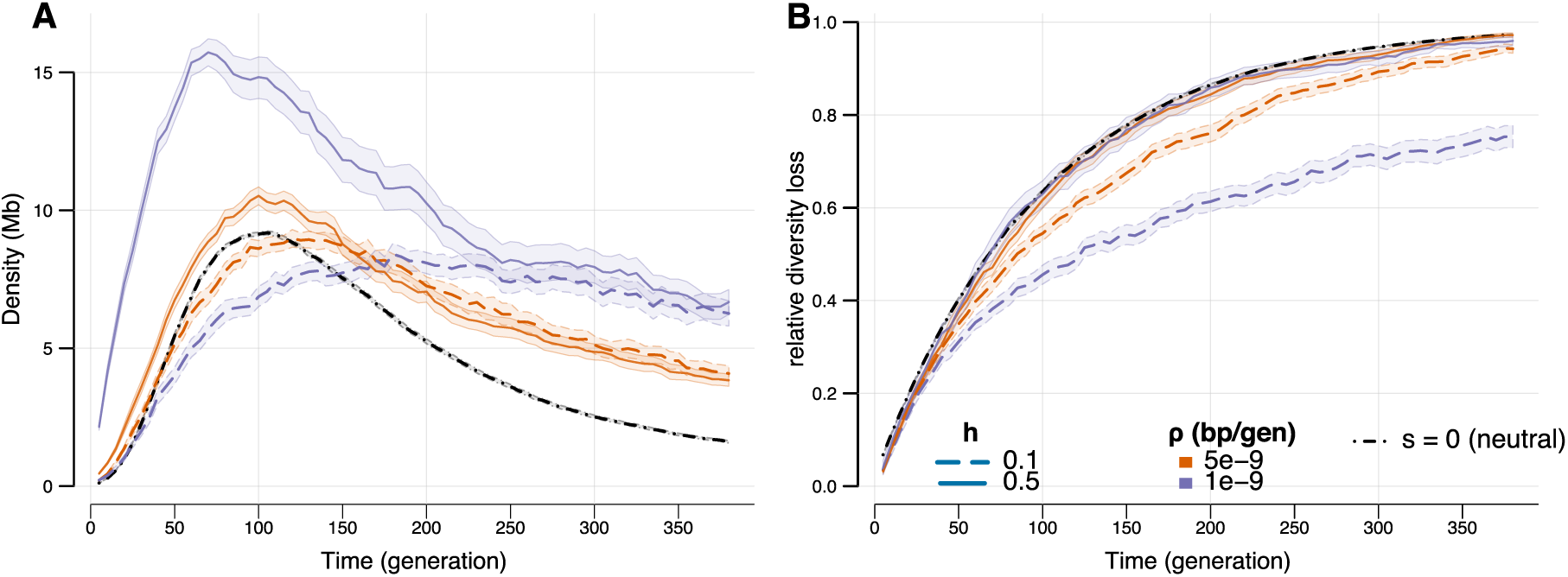
Effect of recombination rate (*ρ*) and dominance levels (*h*) on through density (A) and diversity loss (B) dynamics during a bottleneck. The selection coefficient of deleterious mutation was set to s = -0.0015 for both recessive (*h* = 0.1, dashed lines) and codominant (*h* = 0.5, solid lines) variants. Results for chromosomes with low recombination (*ρ* = 10^-9^ per bp per generation) are shown in purple, whereas those for regions of intermediate recombination (*ρ* = 5 ×10^-9^ per bp per generation) are shown in orange. Shaded areas show 95% CI obtained from 10,000 bootstrap iterations. Note that the neutral model (*s* = 0) is represented in a black dash-dotted line and was carried out in larger chromosomes (100 Mb instead of 20Mb for selected chromosomes), leading to smaller confidence intervals, but with similar average statistics (see **supplementary fig. S3**).

**Figure 4:**
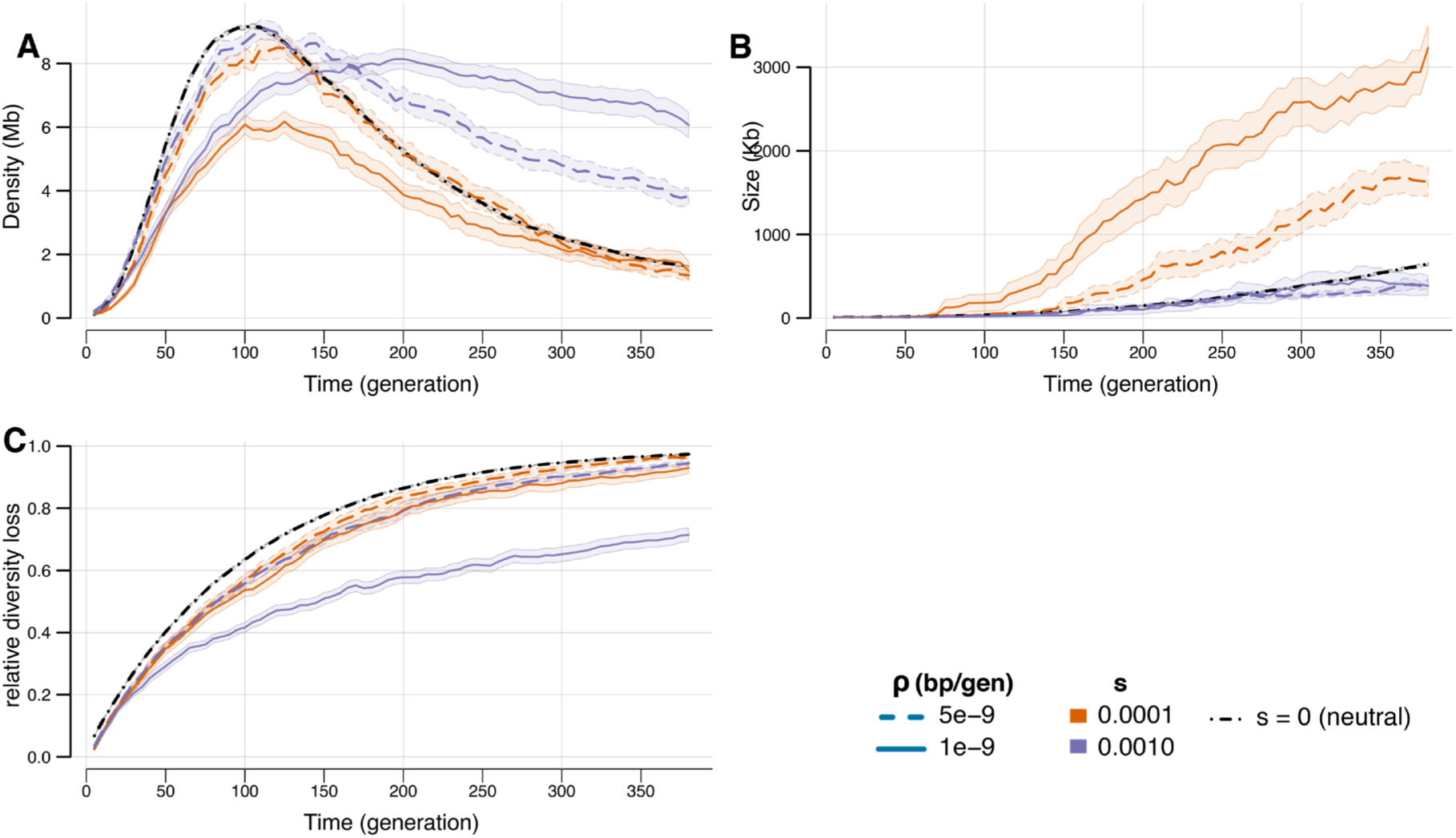
Effect of selection intensity (*s*) on trough density (A), trough size (B) and diversity loss (C) during a bottleneck. All cases shown are for simulations with partially recessive deleterious mutations (*h* = 0.1). Results for chromosomes with a low recombination rate (*ρ* = 10^-9^ per bp per generation) are shown with a solid line, whereas those with an intermediate recombination rate (*ρ* = 5 ×10^-9^ per bp per generation) are shown with a dashed line. Orange lines represent results obtained for chromosomes including deleterious mutation of very small effect (s=-0.0001), whereas those in purple are for chromosomes including mutations 10x stronger (s=-0.001). Shaded areas show 95% CI obtained from 10,000 bootstrap samples.

### Chromosome with recessive deleterious mutations preserve diversity better

In the presence of co-dominant deleterious mutations, trough density on the selected chromosome is always higher than on the neutral chromosome (**fig. 3A**), but the rate of diversity loss is identical for the selected and the neutral chromosome (**fig. 3B**). Therefore, the behaviour of a chromosome with co-dominant mutations is essentially following what is expected from a mere reduction in effective population size by BGS. Contrastingly, compared to a neutral chromosome, one with recessive deleterious mutations shows a delay in trough formation (**fig. 3A**), and importantly, a less rapid loss of diversity during the bottleneck (**fig. 3B**). In other words, initial diversity is better preserved during a bottleneck in regions harbouring recessive mutations than in neutral regions or than in regions including co-dominant mutations, and this is especially true in regions of low recombination (**fig. 3B**). This slower loss of diversity in presence of recessive mutations is illustrated by diversity genome scans done at different time points in **supplementary fig. S2**. This result suggests that pseudo overdominance can also develop during bottlenecks and not only in the ancestral population. This is surprising since one would have expected drift to be stronger than selection during these periods of very small sizes. However, we see that the difference in diversity loss do not immediately emerge, but only after 30 to 50 generations of bottleneck, i.e. after 0.6 *N_Bot_* or 1 *N_Bot_* generations, respectively. Note that chromosomes with dominant deleterious mutations (*h* = 0.8) tend to have a slightly faster rate of diversity loss than those with recessive codominant or neutral mutations (see **supplementary fig. S4).**

### Importance of strength of selection in chromosomes with recessive variants

Since patterns of diversity in chromosomes with co-dominant deleterious mutations can be simply explained by a change in ancestral effective population size, we then focused on chromosomes harbouring recessive mutations associated with different selection coefficients (**fig. 4**). Chromosomes with intermediate recombination rate (*ρ* = 5 ×10^-9^ per bp per generation) harbouring slightly deleterious mutations (*s* = -0.0001) show slightly fewer troughs per Mb than neutral chromosomes (**fig. 4A)**, but these troughs are larger (**fig. 4B)**. Similarly, mutations occurring on chromosomes with lower recombination rates (*ρ* = 1 ×10^-9^ per bp per generation) show much fewer but much larger troughs (**fig. 4A and B)**, such that the overall rate of diversity loss remains close to that of neutral mutations (**fig. 4C**).

Chromosomes with deleterious mutations having a 10 times larger selective disadvantage (*s* = -0.001) show a different pattern. Trough sizes are very similar to those of neutral chromosomes during the bottleneck, but trough density is initially lower than that of neutral chromosomes, but then remains relatively high in the later stages of the bottleneck. As is visible in right pane of **supplementary fig. S2** in generation 200, many troughs can be seen interrupted by very small peaks (islands) of diversity slightly overshooting the trough definition threshold, a pattern not seen at the same time for neutral chromosomes (left pane of **supplementary fig. S2**), explaining the difference in trough density between these cases. Importantly, and as already seen in **fig. 3B**, the rate of diversity loss is much slower in low recombination chromosomes (**fig. 4C**), so that low recombination regions harbouring recessive variants should better preserve diversity during bottlenecks.

Note that the neutral model (*s* = 0) is represented in a black dash-dotted line and was carried out in larger chromosomes (100 Mb instead of 20Mb for selected chromosomes), leading to smaller confidence intervals, but with similar average statistics (see **supplementary fig. S3**).

However, the study of low recombination chromosomes (*ρ* = 10^-9^ per bp per gen.) including recessive mutations (*h* = 0.1) with even larger selection disadvantage (s = -0.01, **supplementary fig. S5**) reveals that, in contrary to what happens in chromosomes with intermediate recombination rates (*ρ* = 5 ×10^-9^ per bp per gen), the rate of diversity loss is not inversely correlated with selective disadvantage, as these chromosomes lose diversity more rapidly than chromosomes harbouring mutations with disadvantage s = -0.001. It therefore seems that in low recombination regions, the presence of mutations with intermediate selection coefficients (s = -0.001) is more efficient in preventing diversity loss than when there are only mutations with either much smaller (s = -0.0001) or much larger (s = -0.01) effect (**supplementary fig. S5**). Note that for chromosomes with codominant variants, the strength of selection has no effect on the rate of diversity loss during bottlenecks, and it has minimal impact on trough density and trough sizes (**supplementary fig. S6**).

### Genomic distribution of deleterious mutations

To better understand what is driving the slower loss of diversity in chromosomes with recessive deleterious mutations, we have examined how these deleterious mutations were distributed across regions of low (troughs) or high (islands) diversity and how this would change over time. The evolution of the mutation load in islands of diversity during bottlenecks is visualized in **fig. 5**. For codominant markers (**fig. 5A**) the genome is invaded by troughs and most genomic windows within islands of diversity harbor no deleterious mutations. In chromosomes with intermediate selected (recessive) mutations and recombination rate (**fig. 5B**), the speed at which troughs invade the genome is slower, and islands of diversity can sometimes maintain 1 deleterious mutation per 10 Kb during the whole bottleneck and very few windows of 10 Kb show 2 or more deleterious mutations. In regions of low recombination with recessive mutations, there is a much higher proportion of the genome in islands of diversity during the bottleneck (**fig. 5C**), explaining the lower rate of diversity loss for this parameter combination (**figs. 3B, 4C)**. Note that we also see here a higher proportion of segments of 10Kb in islands that harbour at least 1 deleterious mutation (**fig. 5C**), and because these islands are relatively large (e.g. around 4Mb at the onset of the bottleneck and 200 and 120Kb after 50 and 100 generations, respectively; see **supplementary fig. S7**), each island contains several sites with deleterious mutations, allowing for the emergence of pseudo-overdominance. With very weakly selected recessive mutations (s = -0.0001), a much larger proportion of the islands of diversity shows 2 or more deleterious mutations (**fig. 5D**), but there are proportionally fewer islands of diversity than for more impacting recessive mutations (s = -0.0015, as in **fig. 5C**). While the evolution of the proportion of the genome in troughs matches the evolution of diversity loss observed in **fig. 3** and **supplementary fig. S5**, we see that the fewer islands of diversity observed in chromosomes with very weakly deleterious recessive mutations show a large excess of multiple deleterious mutations (**fig. 5D**), which is compatible with the emergence of pseudo-overdominance (Glémin 2022).

**Figure 5:**
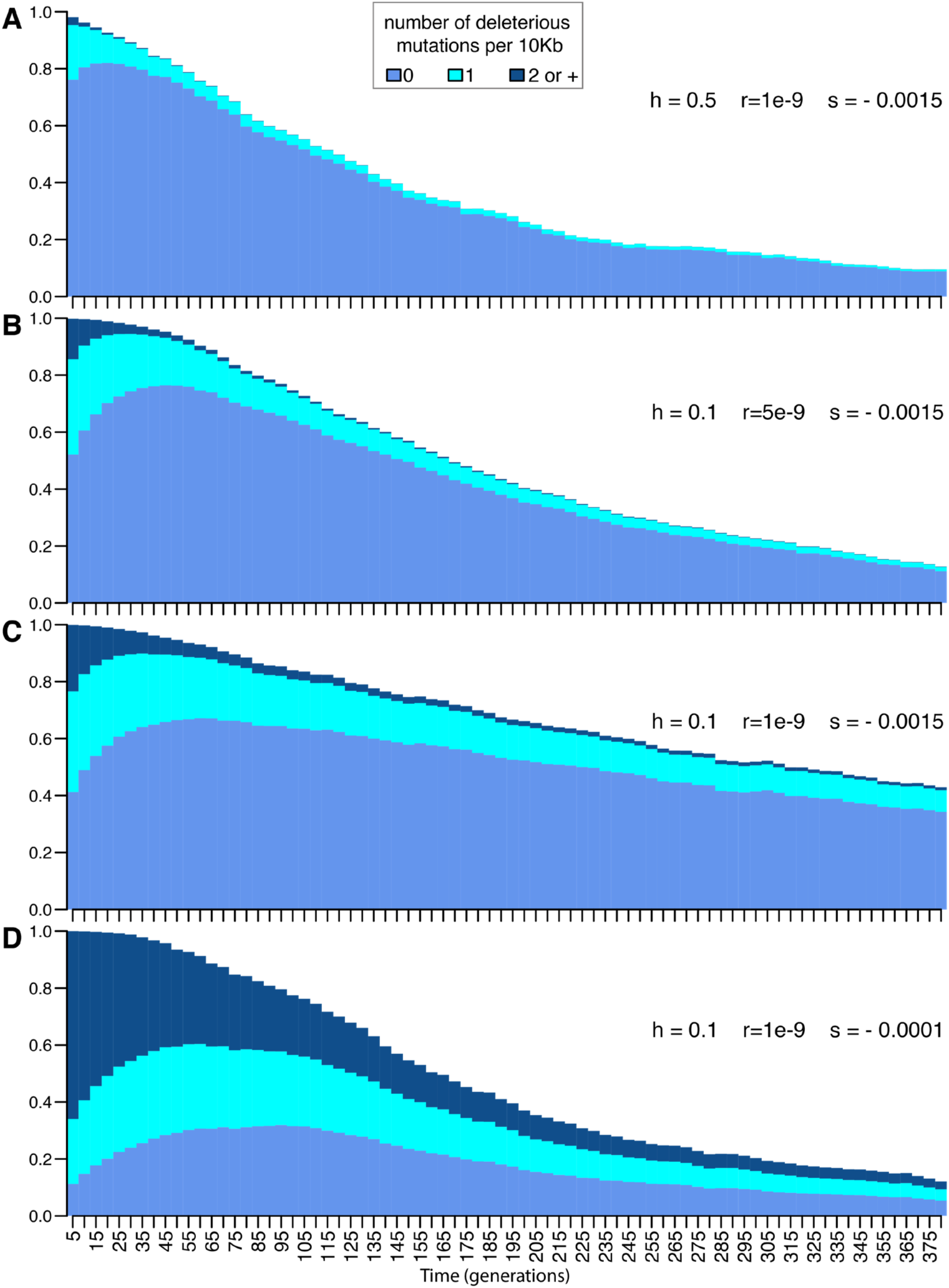
Evolution of the proportion of the genome in islands of diversity (blue colours) including 0, 1, or ≥2 recessive mutations per 10 Kb window. (A) chromosomes of low recombination rate including codominant mutations with selective disadvantage *s* = -0.0015. (B-D) chromosomes with highly recessive mutations (*h* = 0.1). (B) *s* = -0.0015 and intermediate recombination (*ρ* = 5×10^-9^ per bp per generation; (C) same *s* and *h* as B but in regions of low recombination (*ρ* = 10^-9^ per bp per generation); (D) same as C but with recessive deleterious mutations with *s* = -0.0001.

## Discussion

The effect of bottlenecks or range expansions on functional genomic diversity can vary substantially from what is expected in neutral regions as described in Schlichta et al. (2022). Indeed, we find that trough formation is more rapid in regions harbouring co-dominant deleterious mutations than in neutral regions or in regions harbouring recessive mutations (**fig. 3**). Contrastingly, the formation of troughs is slowed down in functional regions with recessive deleterious variants as compared to neutral regions, resulting in a comparatively slower loss of diversity during the bottleneck (**fig. 3**). The preservation of genetic diversity in presence of deleterious variants can only be explained by the emergence of some form of associative overdominance (AOD) (Sved 1968; Ohta and Kimura 1969) and probably of associated pseudo-overdominance (Glémin 2022), where several deleterious alleles are maintained in repulsion on different haplotypes. Whereas AOD has been shown to occur recently in currently large populations (Leitwein et al. 2019; Gilbert et al. 2020), its occurrence in bottlenecked population was unexpected, but it can emerge only for certain combinations of intermediate and small selection coefficients and low recombination rates, the exact combination of which would deserve to be investigated theoretically. Nevertheless, because regions of low recombination are widespread in many species (Stapley et al. 2017) and intermediate selection coefficients of the magnitude used here (e.g. s = ∼0.001) have been inferred in several species (Gallet et al. 2012; Kim et al. 2016; Huber et al. 2017; Kim et al. 2017; Vaughn and Nielsen 2024), conditions for the appearance of AOD during expansions may not be uncommon. The fact that diversity preservation is more pronounced in regions of low recombination suggests that the linkage of multiple low effect mutations in the same chromosomal regions allows selection to be still operational during a bottleneck when the population size is extremely low (*N_Bot_=50)*. This goes against the belief that selection only operates at a given locus if *Ns* is larger than 1 (*Ns* ranged from 0.005 to 0.75 in our simulations) (Kimura 1962), but it is possible in the case of a bottleneck because selection acts at the same time on a large number of selected variants that were created in the ancestral population. Obviously, if the bottleneck was to last for a long time, one would indeed expect the population to reach a new equilibrium, where the fate of new mutations would indeed depend on their associated *Ns* value.

Our work shows that regions of low diversity (troughs) are almost as likely to emerge in functional regions harbouring negatively selected variants than in neutral regions, suggesting that signals looking like selective sweeps are also expected in these regions during bottlenecks or range expansions. In this respect, BGS does not act as a buffering mechanism preventing the emergence of these selective sweep looking regions. In the presence of mainly co-dominant deleterious mutations, one would even expect that these signals be more prevalent than for neutrally evolving regions (**fig. 3**). However, even though most genome will harbour a mixture of dominant, co-dominant and (partially) recessive mutations, it is widely accepted that most functional mutations are partially recessive (Di and Lohmueller 2024), suggesting that results we obtained for recessive mutations should be seen more often, and that troughs will develop more slowly in functional regions than in neutral regions, especially if recombination rates are low. Trough dynamics and genome diversity evolution are also affected by mutation selection intensity (**fig. 4** and **supplementary fig. S5**) and the exact pattern of diversity will depend in a complex way of interactions between recombination rates, dominance levels and selective effects of mutations. Because functional regions with co-dominant mutations lose diversity at the same rate as neutral region, considering that BGS leads to a mere reduction in population size (Charlesworth et al. 1993; Hudson and Kaplan 1995) is a good approximation (**table 1**, **fig. 3B**, **supplementary figs. S4** and **S6**). Contrastingly, functional regions with mainly recessive deleterious mutations should lose genetic diversity more slowly than neutral regions, and this loss is slowest in low recombination regions harbouring partially recessive mutations of intermediate effects (**fig. 4** and **supplementary fig. S5**), results also previously found in Gilbert et al (2020). Note that in the presence of more realistic gamma distributions of fitness effects, the dynamics of trough formation and the rate of diversity loss are relatively close to those of neutral regions, unless the average selection coefficient is large (**supplementary fig. S6**), because most mutations would have very small or very large selection coefficients and would thus have either very little effect or be eliminated very quickly due to their large effect.

While this work initially aimed at studying the trough formation in functional regions during a bottleneck or a range expansion, we believe that the observed difference in loss of diversity between functional and neutral regions is an interesting result. Indeed, the finding that genomic diversity is better preserved in low recombination regions harbouring recessive variants is a testable hypothesis. One could indeed compare neutrally evolving regions with functional regions, since they are expected to show different rates of loss of diversity (**fig. 3**) or compare low recombination with high recombination regions, the latter being virtually unaffected by BGS (Pouyet et al. 2018). However, this comparison might be difficult in practice, since these regions might have different initial levels of diversity (**Table 1**), and functional diversity might always remain lower than neutral diversity during the expansion. Therefore, one would need to have an estimation of the ancestral levels of diversity in those regions, before the start of the bottleneck or range expansion, to be able to make meaningful comparisons.

## Material and Methods

### Forward simulations

We used forward simulations to investigate the effects of recombination and selection on genomic diversity in a diploid population going through a bottleneck (as a surrogate for a population range expansion, see below). We performed all simulations with SLiM 3 (Haller and Messer 2019). First, we simulated the diversity of a stable ancestral population made up of 𝑁_𝑎𝑛𝑐_ diploid individuals (𝑁_𝑎𝑛𝑐_ was set to 10,000 unless specified otherwise) for 10 × 𝑁_𝑎𝑛𝑐_ generations to reach mutation-drift-selection equilibrium. After this burn-in period, we simulated a major instantaneous bottleneck, reducing the population size to *N_Bot_* = 50 individuals, and we let it evolve in isolation for 380 generations (or 7.6 *N_Bot_* generations), an arbitrary time but usually sufficiently long to lose most ancestral diversity. We monitored genetic diversity change by sampling forty diploid genomes every five generations, totalling 76 time-samples for every simulated replicate. We then performed 100 simulation replicates per combination of demographic and selection scenarios.

### Neutral and deleterious mutation rates

For simulations involving deleterious mutations, the genome of each individual was made up of two independent chromosomes of equal length, and neutral diversity was created by using the same neutral mutation rate µ_𝑁𝑒𝑢_ = 1.25e-8 per bp per generation on each chromosome. Note that the value of µ_𝑁𝑒𝑢_is within the range of the average genome-wide rates estimated in humans (Kong et al. 2012; Narasimhan et al. 2017). Background selection (BGS) was simulated on only one of the two chromosomes, by implementing a deleterious mutation rate (µ_𝐷𝑒𝑙_). We set the deleterious mutation rate to 1.1 ×10^-9^ based on the observed reduction of diversity in humans and the theoretical results of Hudson and Kaplan (Hudson and Kaplan 1995). In more details, if the total recombination rate is large relative to the strength of selection against heterozygotes (*sh),* it is predicted that genetic diversity reduced by BGS (𝜋_𝐵𝐺𝑆_) is equal to 𝜋_𝑁𝑒𝑢_ 𝑒𝑥𝑝(− 𝜇_𝐷𝑒𝑙_/𝜌_𝐷𝑒𝑙_) (Hudson and Kaplan 1995), where 𝜋_𝑁𝑒𝑢_is the diversity of neutral regions and 𝜌_𝐷𝑒𝑙_is the per bp recombination rate in the chromosomal region harbouring deleterious mutations. The ratio 𝐵 = 𝜋_𝐵𝐺𝑆_/𝜋_𝑁𝑒𝑢_ is a common measure of the effects of background selection and, when comparing diversity of human African populations in regions with high and low recombination rates taken as proxy for neutral and BGS regions, respectively (Pouyet et al. 2018), *B* is found to be approximately equal to 0.8 (e.g. McVicker et al. 2009). From this observation and under the approximation of Hudson & Kaplan (1995), we infer that μ*_Del_* = −log(*B*) × *ρ_Del_* = 1.1 × 10^-9^, with 𝜌_𝐷𝑒𝑙_ = 5 × 10^-9^ (see below).

In order to test if we can approximate the effect of BGS by a mere reduction of effective population size (Charlesworth et al. 1993; Nordborg et al. 1996; Charlesworth 2013), we also ran simulations with only neutral mutations and investigated the effect of different ancestral population sizes (10000 and 5000 diploid individuals), leading to various levels of ancestral diversity. Note that these pure neutral simulations were done for 100Mb genomes, whilst scenarios with selection were done on a smaller 20Mb genome, for the sake of computational efficiency. We evaluated the effect of genome size on simulated statistics in a few cases (**supplementary fig. S3**) and did not find any significant effect of simulated genome size on trough density and rate of diversity loss, so that all results including selection presented here have been computed from 20Mb genomes.

### Properties of deleterious mutations

Deleterious mutations either had a single fixed selection coefficient, *s*, or selection coefficients were drawn from a distribution of fitness effects (DFE) for each new mutation. For the DFE, we used a Gamma distribution of selection coefficient values around a given mean s and a shape parameter (*a* = 0.23) assumed to fit human functional diversity (Eyre-Walker et al. 2006; Harris and Nielsen 2016). Note that this DFE yields a majority of nearly neutral deleterious mutations and a long tail with higher selective values. The dominance coefficient (*h*) of deleterious mutations in heterozygote individuals was assumed identical for all mutations within replicates such that the fitness of heterozygotes is 1 − ℎ𝑠, whereas the fitness of homozygotes is 1 − 𝑠. We simulated either codominant mutations (*h* = 0.5) or partially recessive mutations (*h* = 0.1). The chosen value of *h* = 0.1 for recessive deleterious mutations is in the range of values previously estimated from humans (0.05 - 0.25; (Agrawal and Whitlock 2011; Di and Lohmueller 2024; Kyriazis and Lohmueller 2024). Individual fitness over the whole genome was then computed multiplicatively across loci for each individual. Finally, the recombination rate was uniform within each chromosome, but different for neutral and selected chromosomes. The recombination rate of the neutral chromosome was set to a “high” rate (1×10^−8^ per bp per generation), whereas that of the selected chromosome was either set to an “intermediate” (5×10^−9^) or “low” (1×10^−9^) rate. These values follow a previous work (Pouyet et al. 2018) showing that the human genome can be safely classified in BGS-affected regions if its recombination rate is smaller than 1cM/Mb or 10^-8^ per bp per generation.

## Summary statistics

Simulated genome sequences were output in VCF format, and then processed by VCFtools (Danecek et al. 2011) and various custom R scripts in written in R 4.3 (R Core Team 2023). The genomes were scanned in 10Kb overlapping windows and expected heterozygosity (𝐻 = 2𝑝(1 − 𝑝), where *p* is the allele frequency) was calculated for each polymorphic locus, summed over all loci, and then divided by the window length, to get the expected nucleotide diversity (𝜋) per bp. As before (Schlichta et al. 2022), we used a threshold of 10% of initial diversity (mean diversity of the ancestral population) to distinguish between troughs and islands of diversity during the bottleneck. Troughs were defined as genomic regions with diversity at or below this 10% threshold, while islands of diversity are regions with diversity above that threshold. Following a previous approach (Schlichta et al. 2022), we summarized the diversity landscape by calculating trough density (number of troughs per Mb), trough size (length) and relative diversity loss (diversity lost compared to the ancestral diversity). For each of the 76 sampling points, all trough statistics were bootstrapped over the 100 replicates to obtain confidence intervals. More specifically, each dataset was resampled with replacement 10,000 times, and statistics were recalculated each time to get the 2.5% and 97.5% quantiles of the distribution, delimiting a 95% confidence interval.

## Data Availability

Scripts used to run simulations and perform data analyses will be publicly available as a git repository (github.com/CMPG/BGS_w_BTNCK).

## Acknowledgements

This work has been supported by a Swiss national Foundation grant No 310030_188883 to LE. We thank the CPMG lab members for constructive discussions during the elaboration of this work.

## Notes

### Competing Interest Statement

The authors have declared no competing interest.

