## supplementary material for "Evolution of functional genomic diversity during a bottleneck"

### Supplementary Materials

*Flávia Schlichta[1,2,*], Stephan Peischl[2,3] and Laurent Excoffier[1,2,*]*

[1] Computational and Molecular Population Genetics (CMPG), Institute of Ecology and Evolution (IEE), University of Bern, 3012 Bern, Switzerland
[2] Swiss Institute of Bioinformatics, 1015 Lausanne, Switzerland
[3] Interfaculty Bioinformatics Unit, University of Bern, 3012 Bern, Switzerland

ORCID:

F. Schlichta: 0000-0002-8845-3623

S. Peischl: 0000-0002-0474-6104

L. Excoffier: 0000-0002-7507-6494

Emails:

F. Schlichta:

S. Peischl:

L. Excoffier:

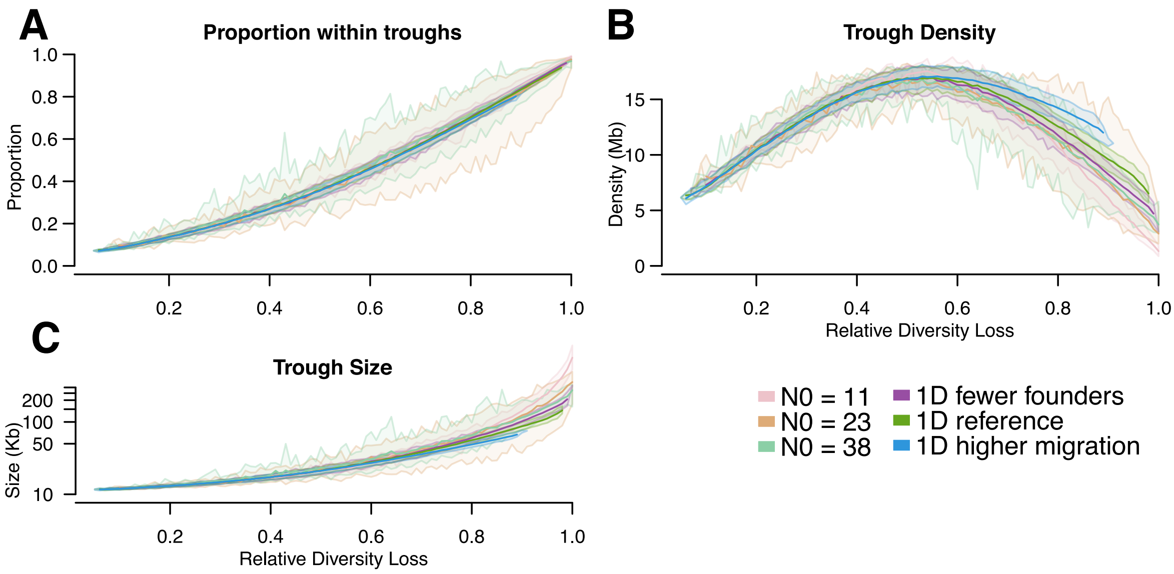


**Figure S1**: Trough formation dynamics during bottlenecks or range expansions starting from the same level of ancestral diversity. A: proportion of the genome in troughs (in regions of the genome with less than 10% of ancestral diversity). B: Number of troughs per Mb. C: Trough size. All trough statistics are plotted as a function of the proportion of diversity lost since the beginning of the bottleneck. We report here trough statistics for bottlenecks of different sizes (N=11, 23 or 38 diploids) and for one-dimensional (1D) range expansions with different properties, either with fewer founders colonizing a new habitat or with higher migration rates between nearby demes, as compared to a reference scenario, as described in Schlichta et al. (2022).

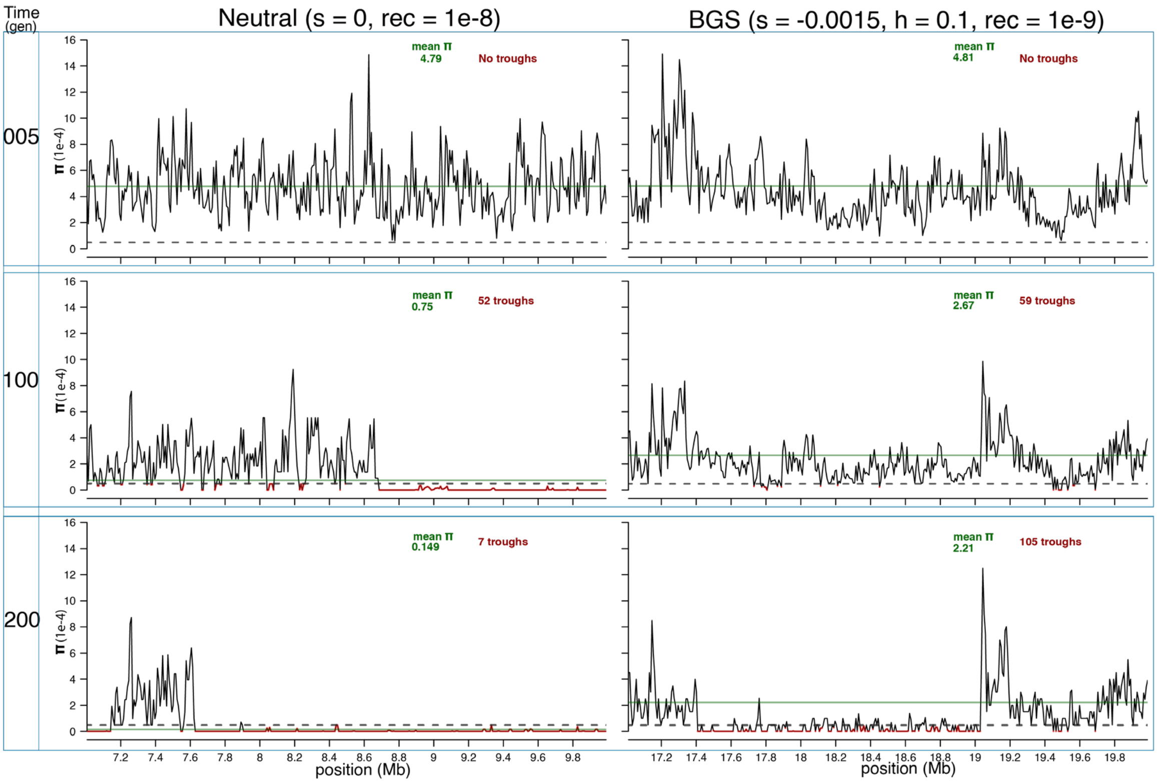


**Figure S2**: Genome scans of nucleotide diversity (π) during a bottleneck, for a neutral (left column) and a selected region (right column, h = 0.1). Three time points are shown: 5, 100 and 200 generations after the start of the bottleneck or a range expansion. Y-axis shows nucleotide diversity and X-axis shows position in the genome (only 3 Mb are shown here for clarity). Horizontal dotted line indicates through threshold (10% of average ancestral diversity within each region); horizontal solid green line shows the average diversity of the chromosome segment at the three-time point (and it is also annotated in green on the top of each panel). Troughs are highlighted in red, with their total number (whole genome) annotated in the top right in red. Note that in this specific comparison, the chromosome with partially recessive deleterious mutations has initially similar levels of diversity than the neutral segment, but it loses diversity less rapidly for a bottleneck of identical intensity.

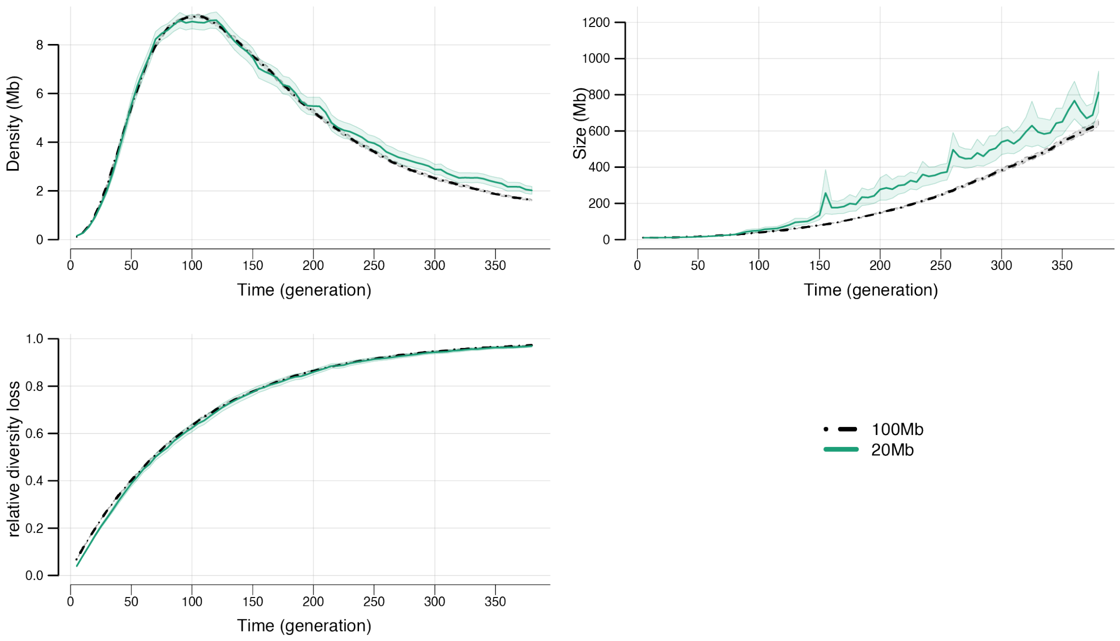


**Figure S3**: Effect of chromosome size on trough formation and genomic diversity during a bottleneck. All mutations are assumed here neutral. Trough density and relative diversity loss in chromosomes of 20 Mb are very similar to what is measured on 100Mb chromosomes. However, trough size is overestimated after 100 generations of bottleneck in chromosomes of 20 Mb.

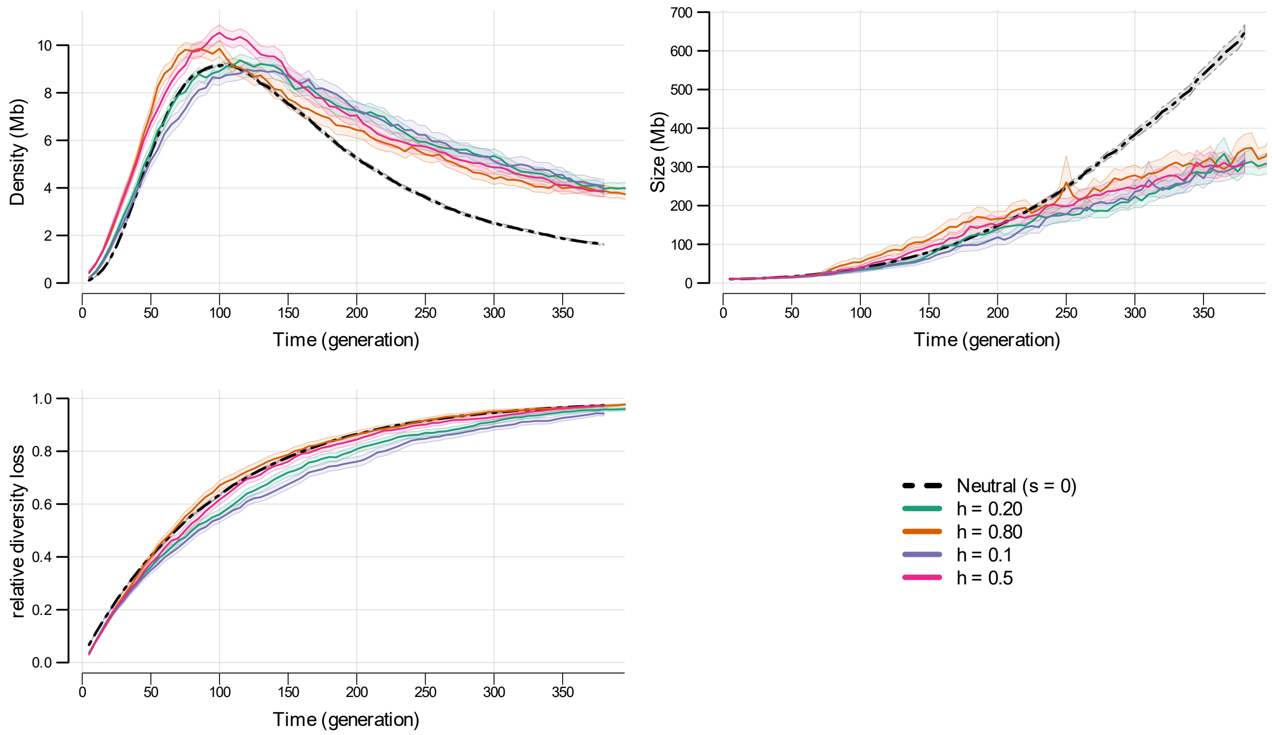


**Figure S4**: Properties of trough formation and genomic diversity loss during a bottleneck for chromosomes harboring mutation with difference dominance levels *h*. We show here simulations done with an intermediate recombination rate of $5 \times 10^{-9}$ per bp per generation.

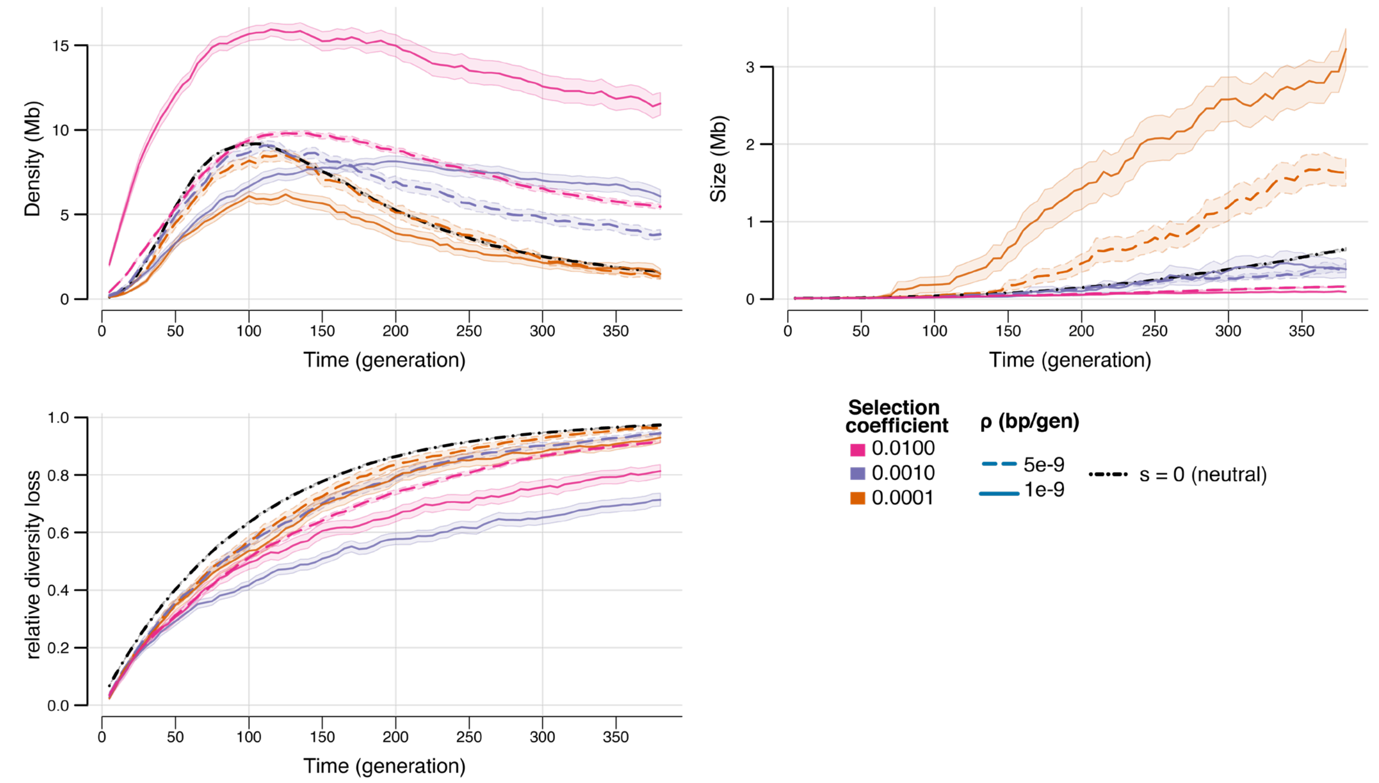


**Figure S5**: Properties of trough formation and genomic diversity loss during a bottleneck for chromosomes harboring recessive mutations (*h*=0.1) and different selection coefficients *s*.

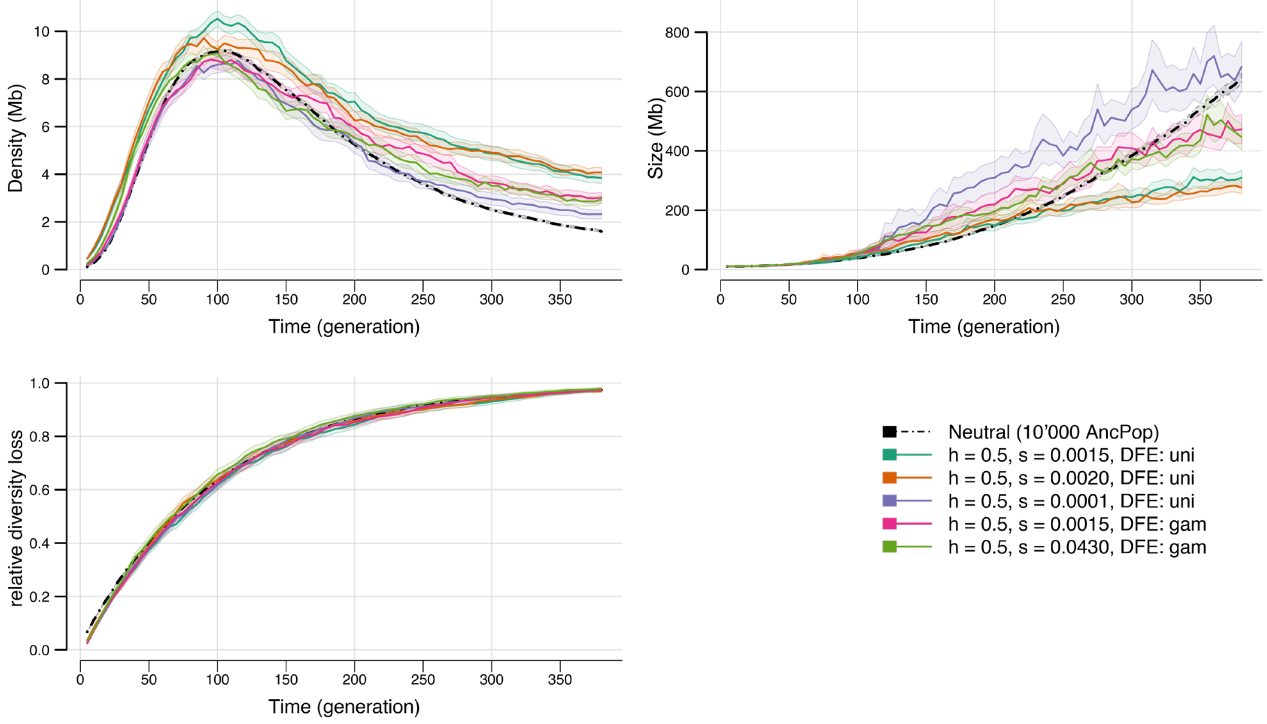


**Figure S6**: Properties of trough formation and genomic diversity loss during a bottleneck for chromosomes harboring codominant mutations (*h*=0.5) and different distributions of selection coefficients, either of constant value, or Gamma distributed value around a certain mean mentioned in the legend pane.

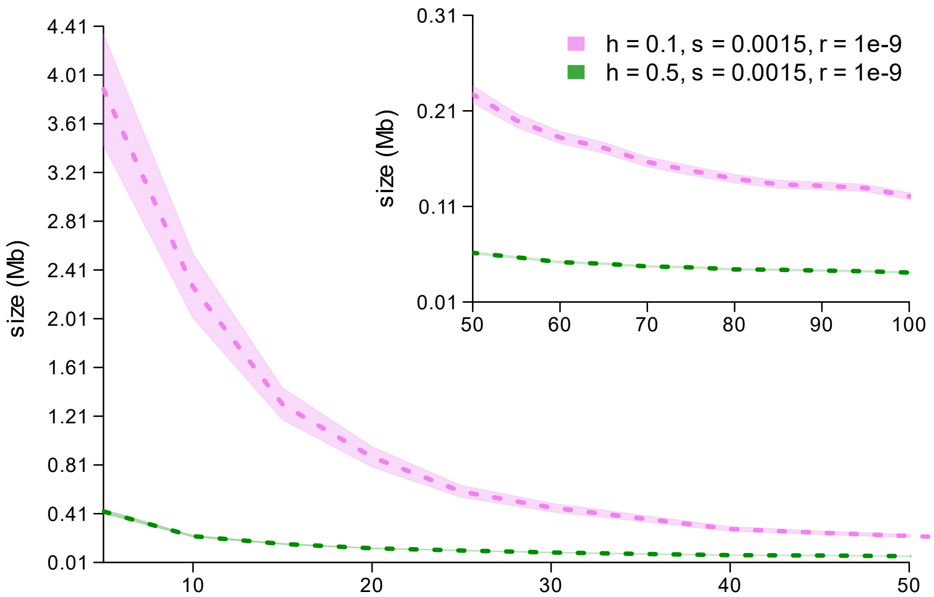


**Figure S7:** Size of diversity islands through time. There is a big difference between having codominant (green) and a recessive (pink) variants in terms of island sizes, and right at the beginning of the bottleneck, codominant islands are 400Kb long compared to ~4 Mb islands formed with highly recessive mutations. There is a quick decrease in island size in both cases, but the recessive mutations still maintain larger islands compared to the codominant variants after 100 generations (~120Kb versus ~40Kb).
